# Single-molecule tracking reveals dual front door/back door inhibition of Cel7A cellulase by its product cellobiose

**DOI:** 10.1101/2023.07.13.548867

**Authors:** Daguan Nong, Zachary K. Haviland, Nerya Zexer, Sarah J. Pfaff, Daniel J. Cosgrove, Ming Tien, Charles T. Anderson, William O. Hancock

## Abstract

Degrading cellulose is a key step in the processing of lignocellulosic biomass into bioethanol. Cellobiose, the disaccharide product of cellulose degradation, has been shown to inhibit cellulase activity, but the mechanisms underlying product inhibition are not clear. We combined single-molecule imaging and biochemical investigations with the goal of revealing the mechanism by which cellobiose inhibits the activity of *Trichoderma reesei* Cel7A, a well-characterized exo-cellulase. We find that cellobiose slows the processive velocity of Cel7A and shortens the distance moved per encounter; effects that can be explained by cellobiose binding to the product release site of the enzyme. Cellobiose also decreases the binding rate of Cel7A to immobilized cellulose but does not slow the binding rate of an isolated carbohydrate-binding module, suggesting that cellobiose inhibits binding of the catalytic domain of Cel7A to cellulose. In support of this, cellopentaose, which is considerably larger than cellobiose, also slows the binding rate of Cel7A to cellulose without affecting the velocity and run length. Together, these results suggest that cellobiose inhibits Cel7A activity both by binding to the ‘back door’ product release site to slow activity and to the ‘front door’ substrate binding tunnel to inhibit interaction with cellulose. These findings point to new strategies for engineering cellulases to reduce product inhibition and enhance cellulose degradation, supporting the growth of a sustainable bioeconomy.

**Significance:** Cellulose, a polymer of repeating glucose subunits, is the primary component of plant cell walls. A promising route to reducing petrochemical use is digesting plant biomass to glucose and fermenting glucose to bioethanol. Cel7A is a model cellulase enzyme that degrades cellulose from one end to generate the disaccharide product, cellobiose. Because industrial-scale bioethanol generation generates high concentrations of cellobiose, product inhibition is a significant concern. We investigated product inhibition of Cel7A by cellobiose at the single-molecule level and found that cellobiose both slows the movement of Cel7 along cellulose and inhibits the initial binding of Cel7 to cellulose. These results suggest that cellobiose binds to the enzyme at more than one site and achieves its inhibition by multiple mechanisms.

## Introduction

Cellulose, the most abundant biopolymer on earth, is a linear polysaccharide consisting of β-1,4-linked D-glucose units arranged in structurally repeating cellobiose units (1) that are released from cellulose during hydrolysis by cellulase enzymes. Because cellulose can be degraded into fermentable sugars for subsequent conversion to renewable fuels and other high-value products, it has enormous potential as a renewable source of energy and biomaterials (2). In nature, degradation of the cellulose polymer into cellobiose is carried out by extracellular cellulase enzymes secreted by fungi and bacteria (3). However, the β-1,4 bonds linking the sugar subunits in each cellulose chain are highly stable (4, 5), and the chains are tightly packed into partially crystalline microfibrils in which only a fraction of the chains lie on the microfibril surface and are thus accessible to enzymatic attack (2). This structure, coupled with the lignin and hemicellulose that surround cellulose in plant cell walls, makes lignocellulosic biomass highly resistant to enzymatic degradation. Intense research efforts are currently focused on improving the hydrolytic degradation of lignocellulose, including advances in biomass pre-treatment technologies and schemes to improve the cellulolytic enzymes that catalyze the conversion of cellulose to fermentable sugars (6-14).

An additional hurdle to cost-effective bioenergy production from plant biomass is that currently employed biomass-degrading enzyme systems are substantially inhibited by hydrolysis products including cellobiose and glucose (11, 14-18). Product inhibition retards the overall conversion rate of cellulose to the final glucose product and is particularly prominent at the high substrate loadings utilized industrially. Among biomass-degrading enzymes, Cel7A derived from the fungus, *Trichoderma reesei (Tr*Cel7A, hereafter Cel7A), is a prominent cellulase that has served as a model enzyme for several decades. Extensive investigations have demonstrated that Cel7A is inhibited by cellobiose (14, 17, 19), which is hypothesized to originate from the high binding affinity of cellobiose for the enzyme’s product release site (20); however, experimental evidence supporting this hypothesis is limited.

Cel7A, which consists of a carbohydrate-binding module (CBM) and a catalytic domain (CD) containing a substrate-binding tunnel, hydrolyzes crystalline cellulose processively from the reducing end (Fig. 1 A) (21). The tunnel encompasses nine glucose subunits, which are numbered -7 to -1 preceding the active site and +1 and +2 in the product release site (Fig. 1A). Two exposed tryptophan residues, W40 and W376, are located at the tunnel’s “front-door” (site -7) and the “back-door” (site +1), respectively. The processive degradation cycle consists of hydrolysis of the β-1,4 glycosidic bond between the -1 and +1 subunits, expulsion of cellobiose from the product release site, and forward movement of the enzyme by two glucose subunits (∼1 nm). Despite extensive study, the mechanism by which cellobiose inhibits Cel7A is not settled. Molecular dynamics simulations suggested a -14.4 kcal/mol free energy for cellobiose binding to the product release site, which corresponds to a 27 pM binding affinity (22). In contrast, functional assays have found half-maximal inhibition at cellobiose concentrations in the 1-20 mM range (23, 24). Numerous models for product inhibition of Cel7A have been put forward. Ståhlberg et al (16) found that added cellobiose had no effect on the adsorption of either the intact enzyme or the isolated carbohydrate-binding module to cellulose, whereas adsorption of the isolated catalytic domain was enhanced (rather than diminished) by cellobiose (16). Lee and Fan (25) suggested the product inhibition mechanism to be the deactivation of the substrate-adsorbed enzyme, a form of uncompetitive inhibition. In contrast, Holtzapple et al. (17) concluded that cellobiose inhibition was noncompetitive and suggested that cellobiose binds to a site that differs from the active site. Finally, Gruno et al. suggested a mixed-type inhibition with an apparent inhibition constant of 1.6 ± 0.5 mM (14).

**Figure 1.**
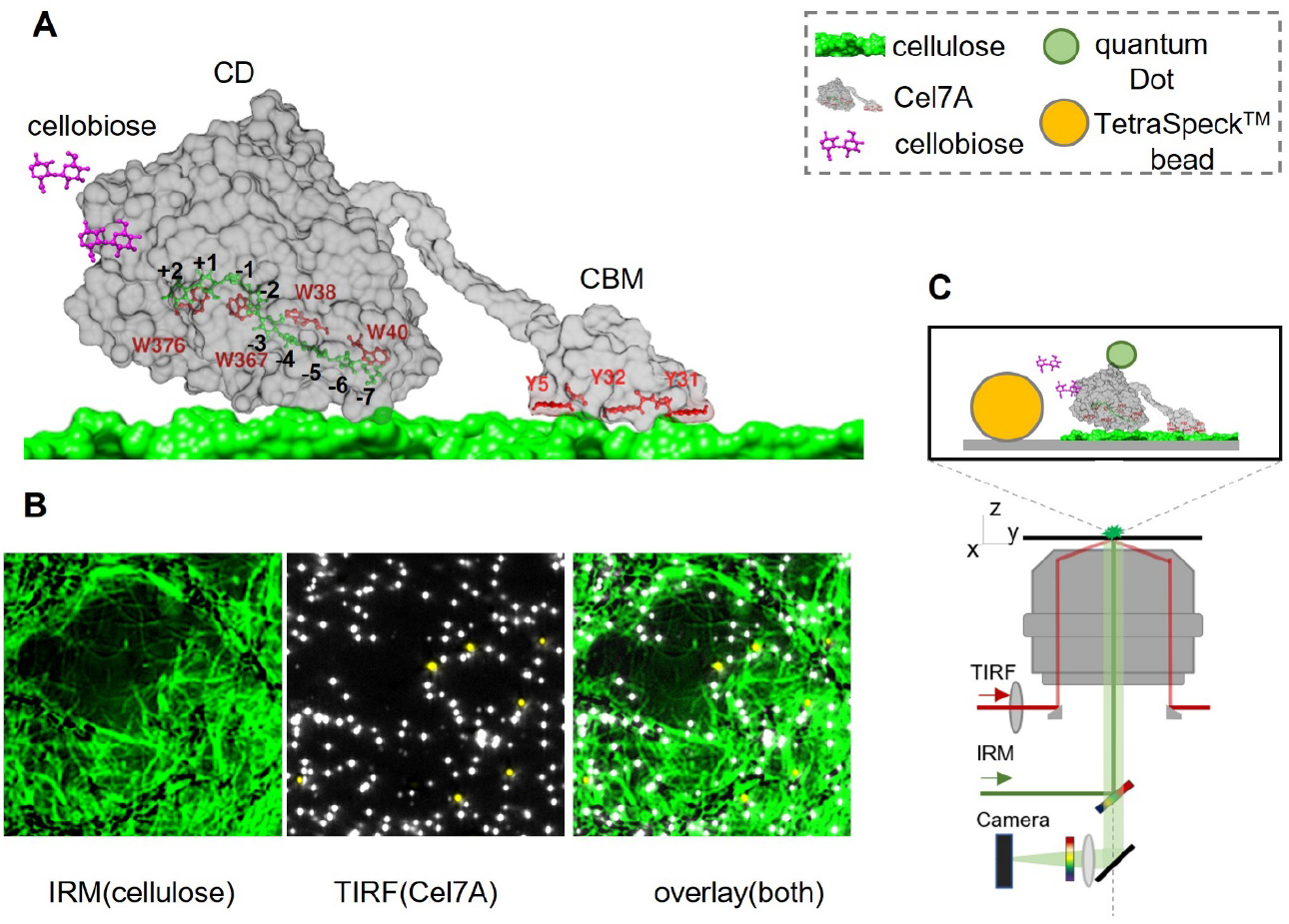
Experimental design. **A)**, Structures of the Cel7A catalytic domain (CD) in complex with a cellulose chain (left, PDB code 8CEL) and the cellulose-binding module (right, PDB code 1CBH). The linker connecting the two domains is drawn by hand. **B)** The surface-immobilized cellulose is imaged by IRM (green strands), Cel7A enzymes (labeled with Qdot525, bright objects) and TetraSpeck beads (yellow dots) are imaged by TIRFM, **C)** Experimental design of Qdot-labeled Cel7 interacting with surface-immobilized cellulose in presence of soluble cellobiose (not to scale); the yellow sphere is TetraSpeck fiduciary marker (29).

The lack of consensus regarding product inhibition of Cel7A is perhaps unsurprising based on the structures of Cel7A and cellulose. In a classical enzyme, the substrate and product binding sites are identical, and product inhibition results from competitive binding to the active site (26). In contrast, in Cel7A cellulose enters through the “front door” of the tunnel and cellobiose is released out the “back door” roughly 5 nm away, with the active site between (Fig. 1A) (20). Furthermore, in addition to the catalytic domain, Cel7A contains a CBM (Fig. 1A) that might facilitate the initial binding of the enzyme to crystalline cellulose, enhance the affinity of the CD for cellulose, or have other functions (27, 28). Thus, among multiple non-exclusive mechanisms, cellobiose might inhibit Cel7A by interacting with the CBM and preventing binding of Cel7A to crystalline cellulose or by promoting dissociation of the CD from cellulose; it might bind to the front door in the catalytic domain to prevent threading of the cellulose chain into the tunnel; or it might bind to the “back door” product release site and thus block threading of cellulose into the active site. In solution studies, these different mechanisms would be expected to have complex effects on *k*_*cat*_ and *K*_*M*_; this complexity might help explain the diverse and conflicting hypotheses regarding product inhibition mechanisms for cellulases in previous work.

In contrast to studies in bulk solution that derive lumped parameters, single-molecule investigations can measure the processive velocity, run length, and other rate constants on individual enzyme molecules and thus provide novel insights into enzyme function. The goal of this study was to dissect the product inhibition mechanism of Cel7A by using single-molecule tracking to quantify the effect of cellobiose on the binding, processive movement, and dissociation of Cel7A from crystalline cellulose. In previous work, we found that Cel7A binds to and moves along cellulose in runs of ∼30 nm at speeds of ∼3 nm/s, corresponding to a hydrolysis rate of ∼3 cellobiose units/s. These processive events were interspersed with numerous immotile episodes lasting tens of seconds, which may be due to the enzyme failing to find an exposed reducing end of a cellulose chain, being unable to extract a cellulose chain to cleave, or other mechanisms. Here, we find that cellobiose not only slows the processive velocity of Cel7A, which is expected, but also slows the landing rate of Cel7A on cellulose. Furthermore, cellobiose diminished the processive run length to a smaller degree than it slowed the velocity, meaning that binding durations were actually longer in the presence of cellobiose. These results suggest a model in which cellobiose inhibits Cel7A both by binding to the product release site to inhibit the forward progress of the enzyme, and also binding to the substrate binding tunnel to inhibit binding of the enzyme to its cellulose substrate.

## Materials and Methods

Isolation of *Gluconacetobacter xylinus* (acetobacter) cellulose and quantum dot (Qdot) labeling of Cel7A (Sigma-Aldrich; Cas: E6412-100UN) were carried out as previously reported (29). Cellobiose was obtained from Sigma (Sigma-Aldrich; Cas: 528-50-7). To investigate the activity of Cel7A on acetobacter cellulose as a function of cellobiose concentration, we adsorbed cellulose to plasma-cleaned glass coverslips by spreading 20 μL of 2.54 mM cellulose on a coverslip, dried it in the oven for 2 min, and assembled a flow cell using double-sided tape. Following cellulose adsorption, surfaces were blocked to minimize nonspecific adsorption by flowing 1 mg/mL bovine serum albumin into the flow cell for 5 min, followed by an enzyme solution consisting of 2 nM Cel7A labeled with 0.5 nM Qdot (Thermo Scientific; Cas: Q10143MP), 5 mM dithiothreitol, and 0 to 16 mM cellobiose in 50 mM sodium acetate buffer, pH 5.0. The enzyme solution was mixed for 5 min before being added to the flow cell. The Qdot525-labeled Cel7A was imaged by a total internal reflection fluorescence microscope (TIRFM) using a 405 nm laser (50 mW) on a custom-built microscope (30). TetraSpeck™ beads (Thermo Scientific; Cas: T7279) were imaged simultaneously as fiduciary markers to compensate for stage drift. Surface-immobilized cellulose was imaged by interference reflection microscopy (IRM), as described (29, 30). Recording of movies began immediately before the enzyme solution was added to the flow cells. Adsorption of Cel7A or CBM3-A488 (31) to the surface of the cellulose (Fig. 2A) was measured by counting bright dots using the “Find Maxima” plugin in ImageJ; prominence was set to 25 for Qdot525-labeled Cel7A and 10 for the Alexa488-labeled CBM3-A488, respectively. Analysis of Cel7A velocity and run length was as described previously (29). All experiments were performed at 21º C.

**Figure 2.**
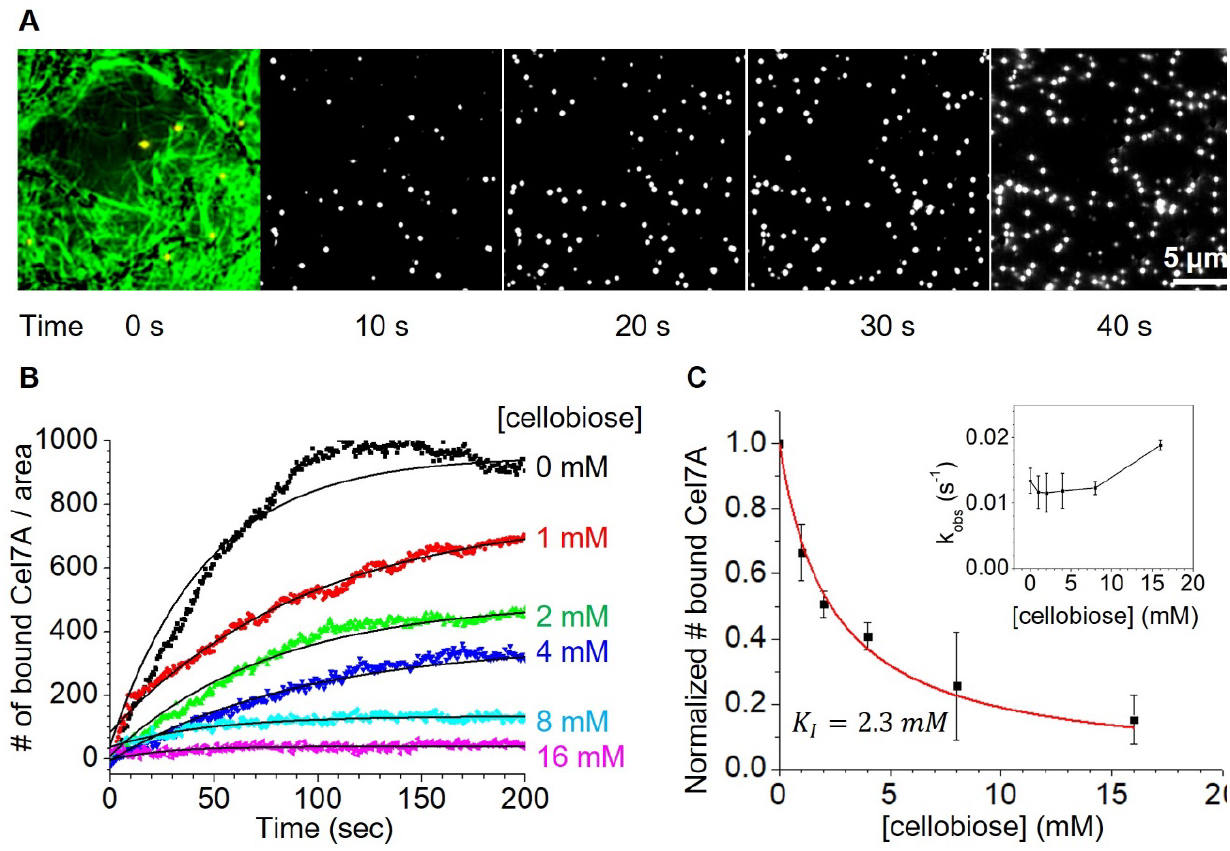
Cellobiose decreases the total numbers of Cel7A enzymes bound to cellulose. Cel7A binding to immobilized cellulose. Image at 0 s shows cellulose substrate (green) imaged by IRM, and TetraSpeck bead fiduciary markers (yellow) imaged by fluorescence. Subsequent images show bound Cel7A (white dots) imaged at 10 s intervals, showing accumulation over time. Numbers of Qdot-labeled Cel7A enzymes bound to surface-immobilized cellulose over time with increasing [cellobiose] from a representative experiment. Values represent numbers of particles per 75 μm by 75 μm field, with values normalized to the proportion of each area covered by cellulose. Binding timecourse data were fit by single exponentials (thin black curves), 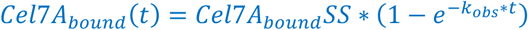. **C)** Steady-state *Cel*7*A*_*bound*_ as a function of [cellobiose]. Data come from three independent datasets, where each is normalized to steady-state *Cel*7*A*_*bound*_ in the absence of cellobiose and error bars represent the standard deviation of three datasets collected on different days. Data are fit by a competitive inhibition model with *Cel*7*A*_*bound*_ = 1/ (1+ [*cellobiose*] /*K*_*I*_), where *K*_*I*_ = 2.3 mM cellobiose. Inset, the observed rate constant *k*_*obs*_ against [cellobiose].

## Results

### Cellobiose decreases the binding affinity of Cel7A to cellulose

To investigate the mechanism of inhibition of Cel7A by cellobiose, we used a previously described single-molecule microscopy assay, in which quantum dot (Qdot)-labeled Cel7A enzymes are visualized landing and moving along bacterial cellulose (29). The surface-immobilized cellulose is imaged by Interference Reflection Microscopy (IRM; Fig. 1B) and the Qdot-labeled Cel7A is imaged by Total Internal Reflection Fluorescence microscopy (TIRF). Upon introducing 2 nM Qdot-labeled Cel7A into a flow cell containing immobilized cellulose, the number of enzymes on the surface increased over time and reached a steady state after roughly 200 seconds (Fig. 2A). We first asked whether cellobiose inhibits the binding of Cel7A with its cellulose substrate. To address this question, we flushed Cel7A into flow cells in the presence of increasing concentrations of cellobiose and monitored the accumulation of bound enzymes on the surface, where the number of surface-bound Cel7A at steady-state reflects a balance of on-rate and off-rate. The steady-state number of bound enzymes decreased progressively with increasing [cellobiose] (Fig. 2B). Each timecourse was fit by a rising exponential function, and the steady-state accumulation as a function of [cellobiose] was well fit by a simple inhibition model with a *K*_*I*_ of 2.3 mM cellobiose (Fig. 2C). The time constant for the rising exponential was independent of [cellobiose] (Fig. 2C inset), a point that we will discuss below.

### Cellobiose decreases the processive velocity and run length of Cel7A on cellulose

We next investigated how cellobiose affects the movement of Cel7A molecules bound to immobilized cellulose. Previously, Cel7A molecules were found to interact with cellulose either in a static state or in a processive manner in which they moved intermittently along the cellulose before stopping or dissociating from the surface (29). These processive movements can be seen in example x-y and distance-time traces of moving Cel7A molecules in the absence or presence of 16 mM cellobiose (Fig. 3A&B). In analyzing processive movements, we defined a moving Cel7A molecule as one that moved at least 10 nm over a duration of at least 5 s, and to avoid false positives due to stage drift, we defined a minimum velocity cutoff of 0.5 nm/s. In the absence of cellobiose, the mean velocity was 4.3 ± 4.9 nm/s (mean ± SD, N = 551 segments) and the run length was 39.7 ± 45 nm (mean ± SD, N = 551 segments), where the removal of each cellobiose unit corresponds to ∼1 nm displacement (32). At 16 mM cellobiose, the distributions of both the velocity and run length were shifted to lower values (Fig. 3C&D): the mean velocity was 1.3 ± 1.7 nm/s (mean ± SD, N = 502 segments) and the run length was 25.1 ± 20.7 nm (mean ± SD, N = 551 segments). When velocity was plotted as a function of [cellobiose], the data were well fit by a simple inhibition model with a *K*_*I*_ of 2.3 mM cellobiose (Fig. 3E). Similarly, Cell7a run length (defined as the distance moved during processive segments) was diminished by cellobiose, with a *K*_*I*_ of 2.6 mM cellobiose (Fig. 3F). However, both inhibition models included an offset, meaning that the velocity was inhibited by a maximum of 75%, whereas run length was inhibited by a maximum of 30%. Thus, cellobiose had a stronger impact on processive velocity than it did on processive run length.

**Figure 3.**
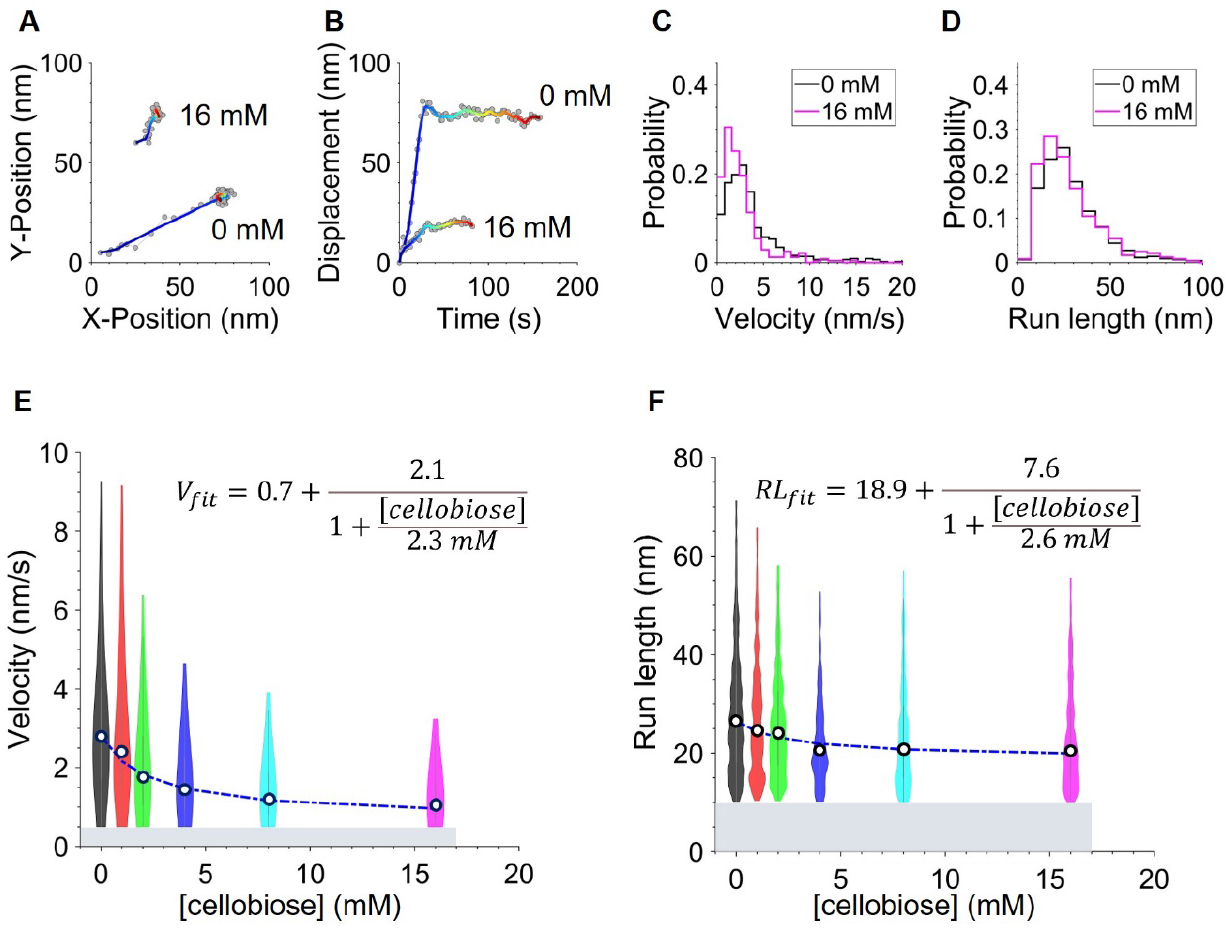
Cellobiose decreases Cel7A processive velocity and run length. **A)** Typical trajectories of Cel7A on the cellulose surface under control and 16 mM cellobiose conditions. Time is color-coded, starting from blue and ending in red. **B)** Distance from origin versus time for the same Cel7A molecules. **C, D)** Distributions of processive velocity **(C)** and run length **(D)** at zero and 16 mM cellobiose concentrations. **E)** Processive velocity of Cel7A as a function of [cellobiose], with fit to a simple inhibition model. Gray bar denotes minimum measurable velocity of 0.5 nm/s. **F)** Processive run length of Cel7A as a function of [cellobiose]. Gray bar denotes minimum measurable run length of 10 nm.

### Cellobiose does not promote dissociation of Cel7A from cellulose

The reduction by cellobiose in the steady-state number of bound Cel7A (Fig. 2) could result from either a reduction in the on-rate for Cel7A binding to cellulose, an increase in the off-rate of the enzyme from cellulose, or both. To address this question, we analyzed the duration of static and processive binding events at increasing cellobiose concentrations. For both static and processive populations, the dwell time that Cel7A was bound to cellulose before dissociation increased slightly with increasing [cellobiose] (Fig. 4A&B). Thus, there is no evidence that the off-rate, defined as the inverse of dwell time, was enhanced by cellobiose. To better understand the effects of cellobiose when the enzyme is actively moving and thus presumably digesting the cellulose, we plotted the duration of processive segments, defined as the run length divided by the velocity. Processive duration increased with increasing [cellobiose] (Fig. 4C). This enhanced duration can be explained by the strong reduction in velocity by cellobiose, (see Fig. 3C) with only a moderate reduction in the run length (see Fig. 3D). Consistent with data for purely static molecules, the duration of static segments that occurred between processive segments for enzymes that moved processively (e.g. Fig. 3A&B) also increased slightly at elevated [cellobiose] (Fig. 4D) (29). Thus, we observed no evidence that cellobiose increases the off-rate of Cel7A from cellulose, and instead the off-rate appeared to slow somewhat with increasing [cellobiose].

**Figure 4.**
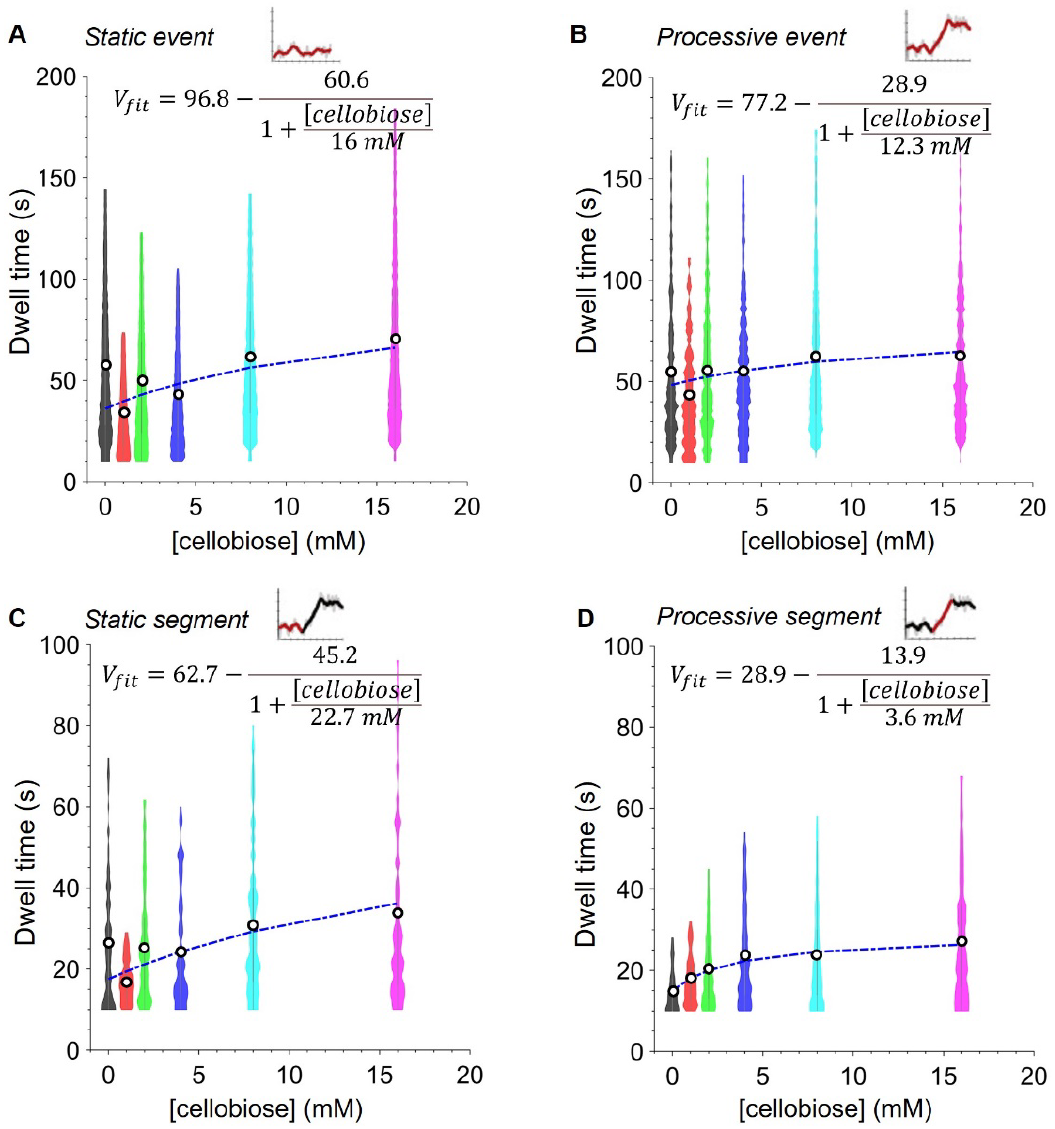
Effect of cellobiose on dwell times of Cel7A in different phases of engagement with cellulose. **A)** Dwell times of Cel7A static binding events, defined as binding events of duration ≥10 s with displacement <10 nm. **B)** Dwell times of processive events, defined as landing events that contain at least one processive segment with displacement >10 nm over at least 10 s. **C)** Dwell times of processive segments, with dwell time calculated as run length/velocity. **D)** Dwell times of static segments that occurred before or after processive segments during processive events. All plots show populations with mean values as open circles. Insets show the distance from origin versus time, with the segments of interest highlighted in red. Curves were fit to a product inhibition model in which cellobiose elongates the dwell times. For panel A, the curve was constrained by *K*_*I*_ ≤ 16 m*M* to allow convergence of the fit; thus, 16 mM is a lower bound.

The lack of an effect of cellobiose on the Cel7A off-rate is consistent with a model in which cellobiose inhibits the binding of Cel7A to cellulose. From Fig. 2, the number of bound Cel7A molecules at steady-state decreased at elevated [cellobiose]. By definition, at steady-state the rate of enzymes binding to the surface is equal to the rate of enzymes leaving the surface. Thus, because the off-rate is unaffected by cellobiose, we conclude that the reduction in the steady-state bound population must result from a decrease in the Cel7A binding rate (developed further in Supplementary Information). These data together suggest that cellobiose acts on Cel7A as a competitive inhibitor of cellulose binding.

### Cellobiose does not affect the binding of a CBM domain to cellulose

Like other cellulases, Cel7A is a modular protein with a N-terminal catalytic domain and a C-terminal carbohydrate binding module (CBM) connected through a polypeptide linker domain (Fig. 1). Thus, Cel7A can potentially bind to cellulose through its catalytic domain, its carbohydrate binding module, or both. As such, cellobiose might inhibit the binding of Cel7A to cellulose by interfering with CBM and/or catalytic domain binding. To test whether cellobiose inhibits binding by a CBM domain, we measured the binding kinetics of CBM3-A488, a recently characterized AlexaFluor488 labled CBM3a fragment from *Clostridium thermocellum* (31). The binding rate was considerably faster than Cel7A, reaching steady-state within roughly 15 s (Fig. 5). More importantly, the steady-state number of bound CBM3-A488 molecules was unaffected by either 50 mM cellobiose or 50 µM cellopentaose. No detectable diffusion of the CBM on the immobilized cellulose was observed. Based on the finding that cellobiose does not affect CBM3-A488 binding to cellulose, we conclude that cellobiose slows Cel7A binding to cellulose by acting on the catalytic domain of the enzyme rather by inhibiting binding through the CBM.

**Figure 5.**
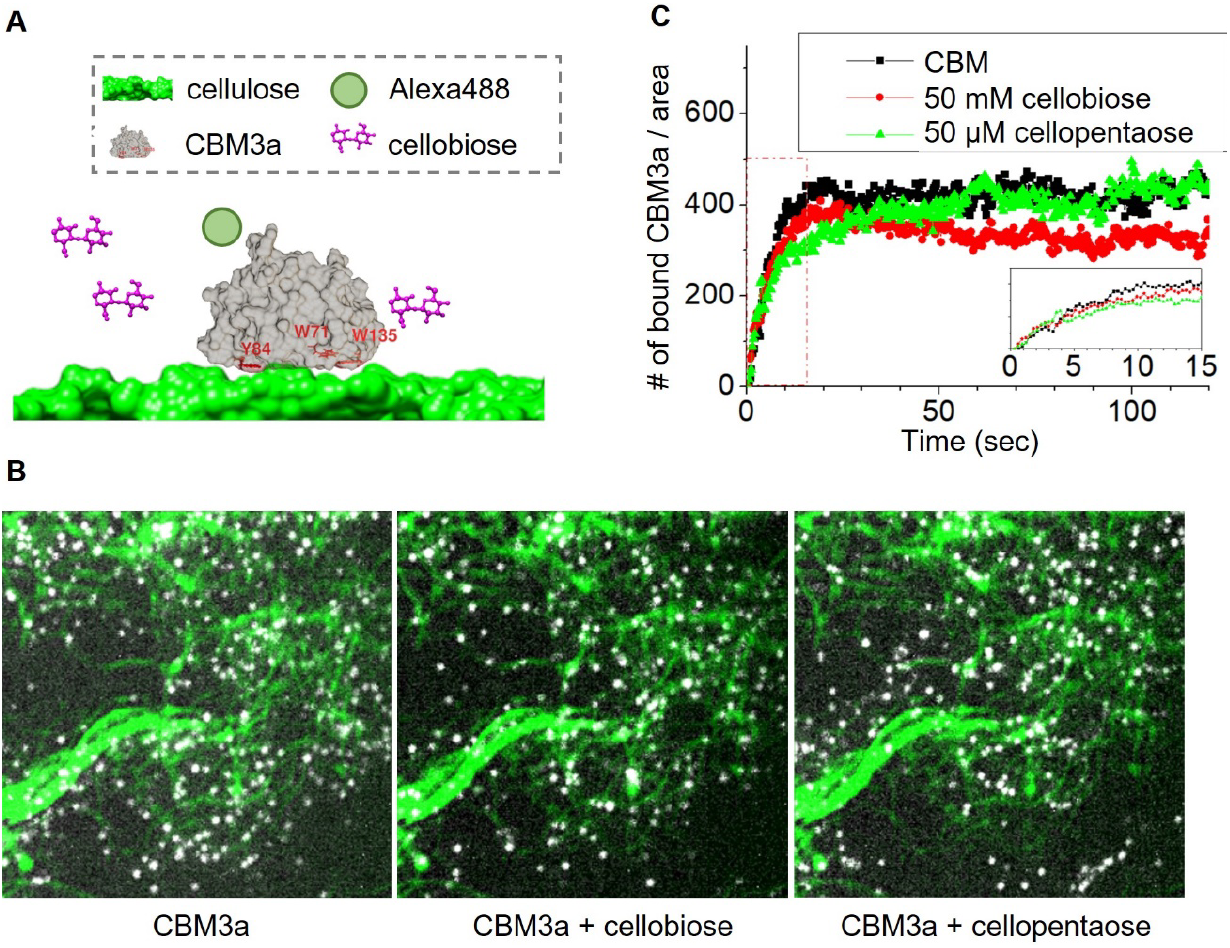
The binding of an isolated CBM to cellulose is unaffected by either cellobiose or cellopentaose. **A)** A model of CBM3a (PDB code: 4JO5) binding to cellulose, showing the location of tryptophan and tyrosine residues implicated in binding. **B)** Steady-state accumulation of 10 pM Alexa488-labeled CBM3-A488 on cellulose under control conditions (left) and in the presence of 50 mM cellobiose (middle) and 50 µM cellopentaose (right). Cellulose imaged by IRM is shown in green, and CBM3-A488 imaged by TIRF is shown in white. **C)** Time course of CBM3-A488 accumulation on cellulose, showing similar landing rate and total number of bound enzymes for the three conditions. Inset shows early landing events.

### Cellopentaose slows the landing rate of Cel7A without affecting its velocity

How might cellobiose inhibit binding of Cel7A to cellulose? One potential mechanism is that cellobiose binds in or near the substrate binding tunnel, the front door, and competitively inhibits the entry of a cellulose chain into the tunnel. If this were the only mode of inhibition (independent of cellobiose binding to the product release site), it should affect the landing rate only and not the velocity. A prediction of this “front door binding” model is that a longer polysaccharide, such as cellopentaose, which can fit into the entrance to the substrate binding tunnel, but which is presumably too large to fit into the product release site, should affect the Cel7A landing rate without affecting the velocity. To test this prediction, we analyzed Cel7A in the presence of increasing concentrations of cellopentaose and repeated the binding rate and enzyme motility analyses we performed for cellobiose. As seen in Fig. 6A&B, cellopentaose inhibited the Cel7A landing rate on cellulose, similar to cellobiose but with a *K*_*I*_ of 1.1 µM, more than 1000-fold tighter than cellobiose. This smaller *K*_*I*_ is consistent with the longer cellopentaose interacting with more residues in the substrate binding tunnel to achieve a higher binding affinity. In contrast with the landing rate, Cel7A velocity and processive run length were unaffected by cellopentaose (Fig. 6C&D).

**Figure 6.**
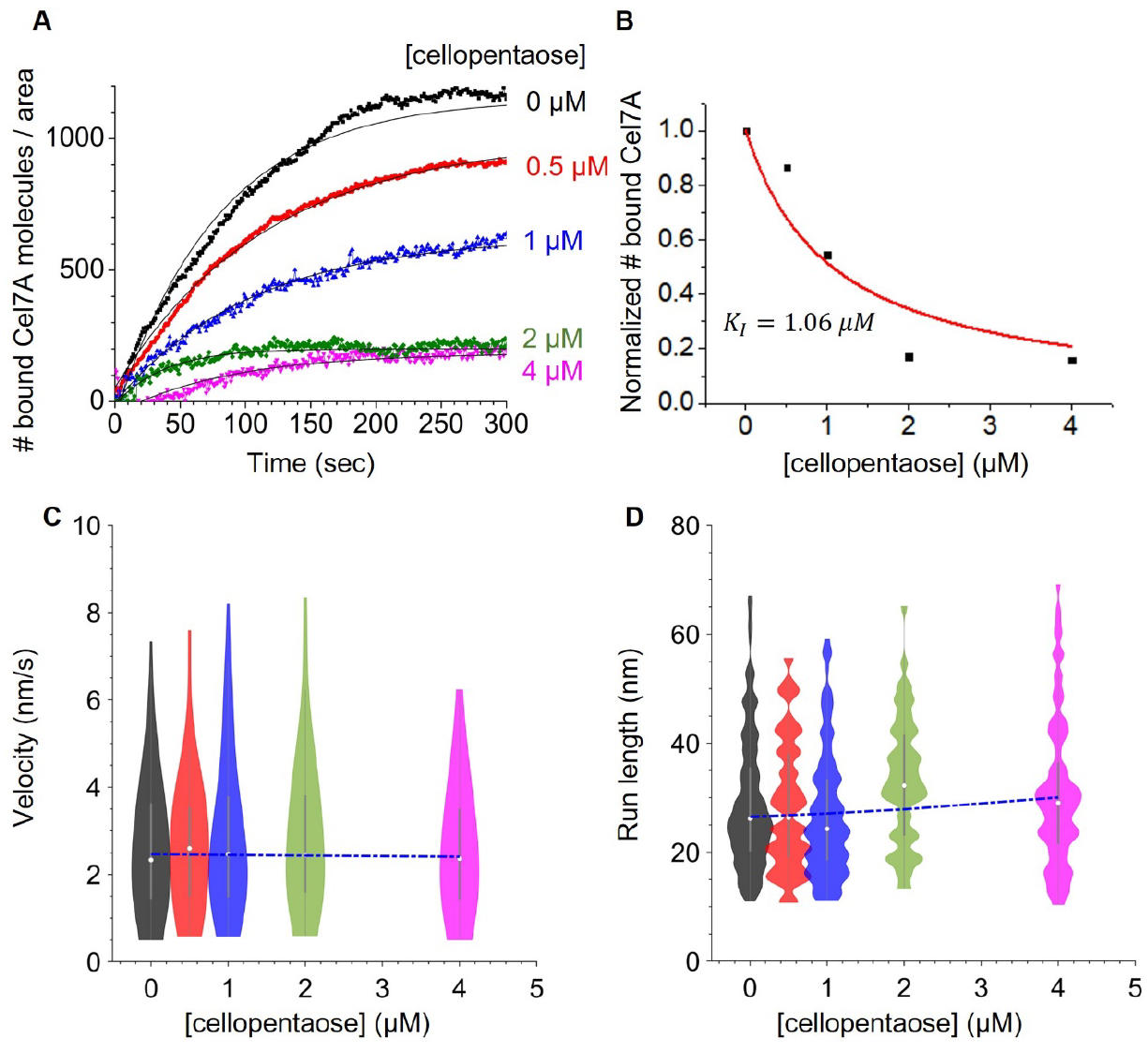
Cellopentaose decreases Cel7A binding to cellulose without affecting the velocity and run length. **A)** Timecourse of the number of Qdot-labeled Cel7A enzymes accumulating on the cellulose surface in the presence of cellopentaose. **B)** Steady-state *Cel*7*A*_*bound*_ as a function of [cellopentaose]. Steady-state *Cel*7*A*_*bound*_ are normalized to control condition in the absence of cellopentaose and are fit by a competitive inhibition model *Cel*7*A*_Bound_ = 1/(1+ [*Cellopentaose*] / *K*_*I*_), where *K*_*I;*_ = 1.1 µM cellopentaose. **(C, D)** Processive velocity and run length as a function of [cellopentaose], showing a lack of inhibition of processive degradation once the enzyme has landed on cellulose.

## Discussion

By quantifying at the single-molecule level how cellobiose alters the landing, processive movement, and dissociation of Cel7A from cellulose, we gain new insights into the mechanism of cellulose degradation by Cel7A. We interpret our results in the context of a model (Fig. 7) in which cellobiose can bind to both the substrate-binding site of Cel7A to inhibit the initial interaction of Cel7A with cellulose, and to the product-release site to block the forward progress of the enzyme.

**Figure 7.**
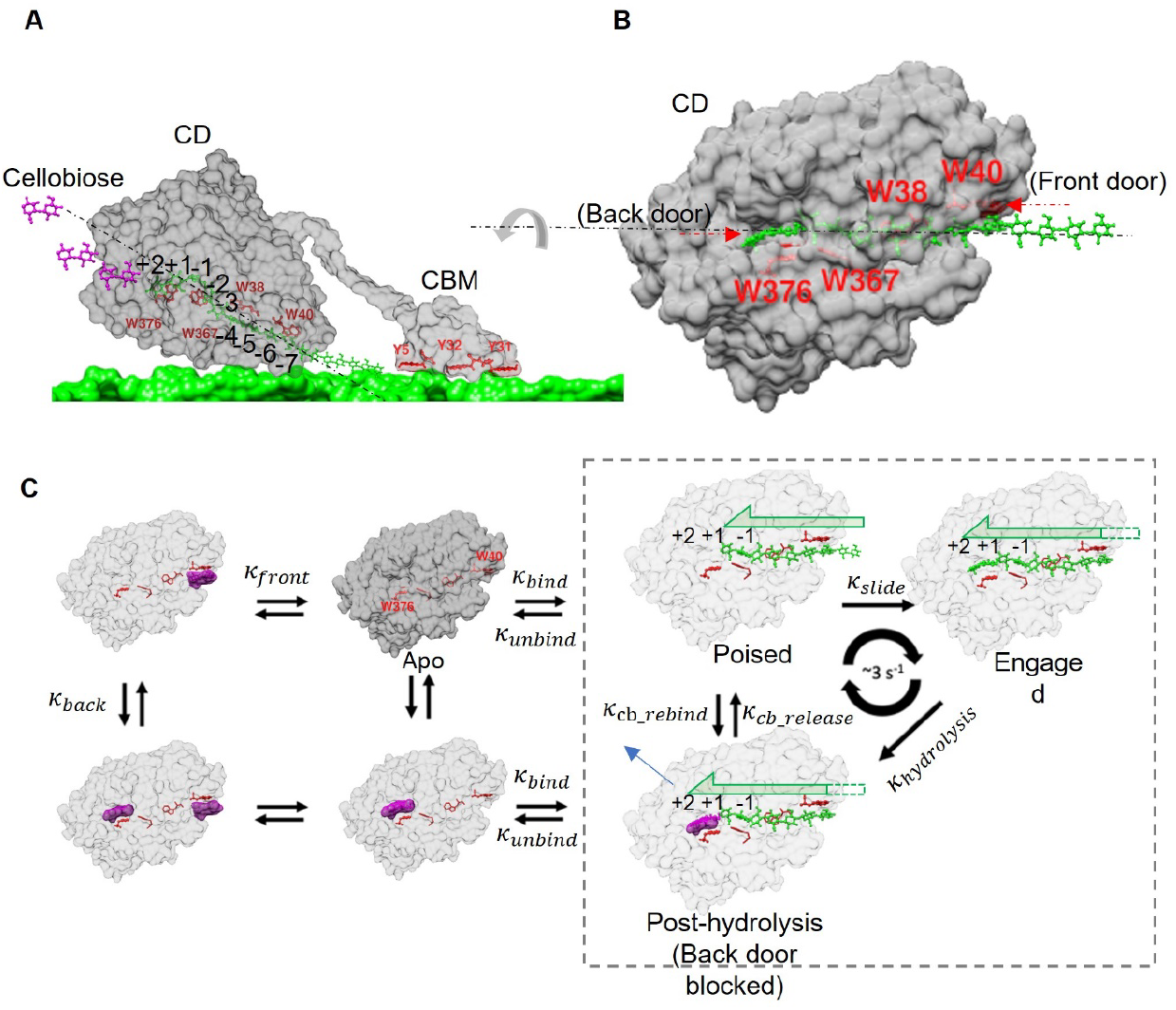
Model of Cel7A inhibition by cellobiose. A) Structures of the Cel7A catalytic domain (CD; PDB code 8CEL) in complex with a cellulose chain, the cellulose-binding module (CBM; PDB code 1CBH) adsorbed to the crystalline cellulose lattice, and cellobiose (purple) being released from the product binding site. B) The catalytic domain is rotated to show entrance of the substrate binding tunnel at the -7 site. C) Model of Cel7A inhibition by cellobiose. Free Cel7A in solution (Apo state) binds to an exposed reducing end of a cellulose strand with rate *k*_*bind*_ to enter the Poised state in which the product release site is empty. Cel7A slides forward at rate *k*_*engage*_ to enter the Engaged state in which the strand is positioned in the active site of the enzyme. Hydrolysis of the cellulose strand at rate *k*_*hydrolysis*_ generates cellobiose in the active site, and cellobiose is released at rate *k*_*cb_release*_ to complete the processive cycle. This processive cycle occurs at ∼3 s^-1^ and results in a 1 nm displacement of the enzyme. Cellobiose (purple) can inhibit binding of Cel7A to cellulose by binding to the front door of the enzyme (*k*_*front*_); the apparent binding constant, *K*_*I*_ is 2.3 mM based on Fig. 2C. Cellobiose can slow the catalytic cycle by binding to the product release site of Cel7A in the Poised state (*k*_*cb_rebind*_) thus inhibiting forward sliding; the *K*_*I*_ for slowing the catalytic cycle is 2.3 mM based on Fig. 3E.

A unique aspect of processive glucoside hydrolases like Cel7A is that their substrate, a cellulose chain, is threaded through the enzyme to the active site, thus separating the sites of substrate binding and product release. Enzymes that release products through a different route than they bind their substrates have been termed “back door” enzymes, with examples including acetylcholinesterase, myosin and actin (33-35). By this definition, Cel7A is a back door enzyme; the substrate binding channel is the front door, with tryptophan W40 being a key mediator of substrate binding (Fig. 1A) (20), and the product release site is the back door, where tryptophan W376 has been proposed as a key mediator of cellobiose binding (Fig. 1A) (8, 20). This front door/back door structure makes it difficult to infer the mechanism of product inhibition from bulk solution studies alone, and emphasizes the need for single-molecule approaches to uncover the specific steps in the enzymatic cycle that are altered by cellobiose.

We propose that cellobiose slows the velocity Cel7A by reversibly binding to the product release site and inhibiting the forward movement of the enzyme along cellulose (Fig. 7). Because Cel7A is processive and remains bound to cellulose before and after product release, inhibition of its velocity by cellobiose is expected to be non-competitive. The 2 mM *K*_*I*_ for inhibition of velocity agrees with previous results from bulk solution studies (23, 24), although it diverges from Molecular Dynamics simulations that predict the affinity of cellobiose for the product release site to be 28 pM (based on a ΔG of -14.4 kcal/mol (22)), and isothermal calorimetry experiments that measured a 19 µM affinity for cellobiose binding to *Talaromyces emersonii* Cel7A (11). How can we reconcile a micromolar cellobiose affinity in solution with a mM observed inhibition constant for processive velocity? Focusing on the Poised state in Fig. 7, one possibility is that the presence of a cellulose chain in the substrate tunnel allosterically lowers the affinity of the product release site for cellobiose, and this conformation is not accessed in the published experiments and simulations of cellobiose binding to Cel7A. An alternate explanation is that progression of the enzyme along the cellulose chain from the Poised state to the Engaged state is strongly favored kinetically and/or thermodynamically over binding cellobiose from solution, which returns the enzyme to the Post-hydrolysis state.

It is notable that the Cel7A dissociation rate was not accelerated at high [cellobiose] where the enzyme was slowed considerably (Fig. 4). One outstanding question regarding the processive mechanism of Cel7A is: *What terminates a processive run of Cel7A?* Previously, we measured a mean run length of ∼30 nm for single Cel7A molecules on immobilized bacterial cellulose (29). This value is approximately an order of magnitude shorter than estimates of average cellulose chain length of ∼300 cellobiose equivalents (36), implying that processive runs are not terminated by the enzyme reaching the end of a strand. One potential termination mechanism is the cellulose chain dethreading (37) out of the tunnel, shown as dissociation from either the Poised or Post-hydrolysis states in Fig. 7. One prediction from this mechanism that is contradicted by the data reported here is that slowing the processive cycle by cellobiose should speed dissociation of Cel7A. Specifically, if hydrolysis were the rate-limiting step in the processive cycle (21, 38), and if dissociation does not occur from the Engaged state in Fig. 7 (for instance, if fully engaging the strand to occupy the +1 and +2 sites increases the binding affinity), then in the absence of cellobiose, the enzyme will predominantly reside in the Engaged state, whereas at high [cellobiose], the enzyme will predominantly reside in the Post-hydrolysis state. One mechanism to account for the finding that the processive dwell time increases, rather than decreases in the presence of cellobiose (Fig. 4C) is that Cel7A dissociates only from the Poised state and not from the Post-hydrolysis state. In this way, cellobiose acts as an uncompetitive inhibitor in that it enhances binding of the enzyme to the cellulose substrate by slowing dissociation from the cellulose substrate.

One unexpected finding was that the binding of Cel7A to cellulose was diminished at increasing cellobiose (and cellopentaose) concentrations. Binding of Cel7A to crystalline cellulose involves a multi-step process of binding to the cellulose surface (through either the catalytic domain or carbohydrate binding module), finding a free reducing end, and the enzyme threading the chain into the substrate tunnel to fully engage with the substrate. This binding process is simplified into the single substrate binding step in Fig. 7. The slowing of Cel7A binding kinetics by cellobiose can be seen by comparing the landing rate curves to the dwell time curves. With elevated [cellobiose], the dwell time did not shorten, and instead was slightly longer (Fig. 4), which can be explained by a substantial slowing of the Cel7A processive velocity coupled with a weaker effect on the run length. Thus, the Cel7A off-rate, which is calculated by inverting the dwell time, was not enhanced at elevated [cellobiose]. Turning to the landing rate data in Fig. 2, we can see that the steady-state population of bound Cel7A decreased strongly with increasing [cellobiose], whereas the observed rate constant describing the exponential rise to the steady-state plateau did not change. The decrease in the plateau without any change in the rate constant of accumulation is surprising, because in a standard two-component equilibrium, the exponential rate constant that describes the response of a perturbation (flushing the Cel7A into the flow cell in this case) generally involves both the forward and reverse rate constants (39). However, as described in Supplementary Information, both the cellulose binding sites and the number of Cel7A enzymes are in excess under the conditions of the experiment. In this special case, the steady-state plateau is proportional to *k*_*on*_ ∗ [*Cel*7*A*] /(*k*_*on*_ ∗ [*Cel*7*A*] + *k*_*off*_), whereas the accumulation rate is determined solely by the off-rate, *k*_*on*_. From this analysis, we conclude that cellobiose affects only the on-rate of Cel7A for cellulose and not the off-rate.

How does the cellobiose product inhibit Cel7A binding to its substrate, cellulose? The first mechanism we considered is that binding of Cel7A to crystalline cellulose occurs via the CBM, and cellobiose binds to the same site on the CBM and competitively inhibits cellulose binding. We collected evidence against this model by finding that cellobiose has no effect on the binding dynamics of an isolated CBM to cellulose (Fig. 5). Two caveats of this experiment are that the CBM is from a different cellulase and that the observed binding rates were considerably faster than for intact Cel7A. Nevertheless, this result is consistent with previous work showing that Cel7A is fully functional in the absence of its CBM (6, 40, 41). A second mechanism we considered was that cellobiose binds to the product release site (the back door), which blocks Cel7A binding to crystalline cellulose. However, because these two sites are on opposite sides of the catalytic domain and engagement with a cellulose strand involves sites -7 to -1, it is difficult to explain, based on our current understanding, how cellobiose could affect the Cel7A on-rate by binding to the back door. This ‘back-door-only’ model is also contradicted by the finding that cellopentaose, which is too large to fit into the product release site (20), also inhibited the landing rate while having no effect on velocity (Fig. 6). Thus, we propose that cellobiose acts as a competitive inhibitor of cellulose by binding to the substrate binding tunnel of Cel7A (the front door) and preventing the enzyme from productively binding to cellulose (Fig. 7). Our data are also compatible with a model in which Cel7A binding to cellulose is a two-step process involving an initial adsorption step followed by threading of the cellulose strand into the tunnel; in this case, it is this initial adsorption step that is inhibited by cellobiose.

Our results suggest that the disparate conclusions in the literature regarding the mechanism of product inhibition of Cel7A arise from two factors: first, Cel7A is a processive enzyme that acts on an insoluble substrate and thus likely differs from classical models of enzyme inhibition, and second, cellobiose binds to at least two separate sites on the enzyme. Cellobiose binding to the front door of the substrate binding tunnel is a form of competitive inhibition, which is expected to raise the *K*_*M*_ for cellulose and have no effect on the *k*_*cat*_. In contrast, cellobiose binding to the product release site is expected to slow the *k*_*cat*_, which in its simplest form is noncompetitive inhibition. However, because cellobiose slows Cel7A velocity to a greater degree than it decreases run length (Fig. 3E & F), the Cel7A off-rate is slowed somewhat by cellobiose, which is expected to decrease the *K*_*M*_, a hallmark of uncompetitive inhibition. Using the steady-state accumulation data in Fig. 2C together with the velocity data in Fig. 3E, we simulated expected results from a bulk biochemical assay at varying [cellobiose]. We found that the simulated data fit a mixed inhibition model, with the *k*_*cat*_ decreasing and *K*_*M*_ increasing with increasing [cellobiose] (Supplementary Data).

How can these new insights into Cel7A product inhibition help efforts to more cost-efficiently convert lignocellulosic biomass to bioethanol? One clear direction is to explore engineered Cel7A with mutations in the substrate binding tunnel and product release site that reduce the affinity for cellobiose without inhibiting binding or hydrolysis of cellulose. To that end, mutating W40 in the front door of the substrate binding tunnel was found to increase the *K*_*M*_ for cellulose by a factor of two, while also having the added benefit of increasing the *k*_*cat*_ (6, 42). A promising avenue for future work will be exploring the degree to which product inhibition is diminished in this and other mutations located in the substrate tunnel. In principle, an even more promising direction is to mutate residues around the product release site to reduce cellobiose affinity at the back door. However, a published study that explored a large number of back door mutants found that, although some mutations (including the equivalent of W376A; Fig. 1A) did reduce the extent of product inhibition, they also all diminished the overall turnover rate of the enzyme (11). One implication of the current results is that mutations at the front door are expected to alter the effect of cellobiose on the *K*_*M*_ of Cel7A for cellulose, whereas mutations to the back door are expected to alter the effect of cellobiose on the *k*_*cat*_. A final implication of the current work is that product inhibition does not appear to act through the carbohydrate binding domain, and as such, further engineering of that domain appears less promising than the catalytic domain. Together, these results point to new directions for engineering cellulases as critical components of lignocellulose processing in a sustainable bioeconomy.

## Acknowledgements

This work was supported by the Department of Energy Office of Science grant number DE-SC0019065. Preparation of CBM3-488 was supported as part of the Center for Lignocellulose Structure and Formation, an Energy Frontier Research Center funded by the US Department of Energy, Office of Science, Basic Energy Sciences under award DE-SC0001090.

## Supplementary Information

### Model for Cel7A binding to immobilized cellulose

To analyze the Cel7A binding kinetics to immobilized cellulose, we developed a simple kinetic model as follows.

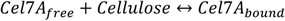

Based on this model:

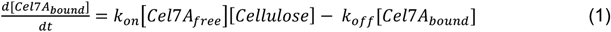

If both Cel7A and cellulose are large relative to the number of Cel7A bound to the surface, then this simplifies the equations. This condition is equivalent to saying there is no depletion of either [Cel7A_free_] or [cellulose] due increasing [Cel7A_bound_] over time, which can be justified as follows.

Cel7A binding in Fig. 2 is shown as number of Qdot-labeled Cel7A molecules per field of view, normalized to the fraction of the field of view taken up by immobilized cellulose. The field of view of our camera is 1200×1200 pixels at 73 nm/pixel, which comes out to 7674 μm^2^. From Fig. 2A, using a 0.5 nM concentration of Qdots in solution, the maximum steady-state accumulation was ∼1000 Qdots per screen. This corresponds to a density of 0.13 Qdot per μm^2^. To determine whether this degree of binding will deplete the Qdot-labeled Cel7A from solution, we consider a 1 μm^2^ area of the surface and the corresponding volume above it in the ∼100 μm thick flow cell. The corresponding volume is 100 μm^3^ = 10^−13^ L. Using Avogadro’s number, a 0.5 nM Qdot concentration in this volume of solution contains (5 × 10^−10^ mol/L) *(10^−13^ L) *(6 × 10^23^ particles/mol) = 30 particles. Binding of 0.13 Qdot per μm^2^ corresponds to <0.5% depletion. Thus, we can make the assumption that despite Qdot-labeled Cel7A binding to the surface, the solution concentration of Qdots remains approximately constant.

Cellulose is adsorbed to the cover glass surface by spreading 20 μL of 2.54 mM cellulose stock solution (expressed as concentration of glucose subunits) over roughly a 1 cm^2^ area of a coverslip. Assuming that all of the cellulose is adsorbed to the surface, this comes out to 5 × 10^−8^ mol of cellulose spread over 10^8^ μm^2^ surface area, for a surface density of 5 × 10^−16^ mol/μm^2^ or 3 × 10^8^ glucose molecules/ μm^2^. We previously measured our bacterial cellulose to consist of one reducing end per 300 glucose subunits (1). This means that in a 1 μm^2^ area on the coverslip surface, there are 10^6^ reducing ends. As described above, the maximum steady state density is 0.13 Qdots/μm^2^. Thus, as long as at least one in every 10^5^ reducing ends are exposed, then there will be negligible (<2%) depletion of cellulose reducing ends by bound Cel7A.

Based on these analyses, we conclude that in our assays, binding of Qdot-labeled Cel7A to surface-immobilized cellulose depletes neither the Qdot concentration in solution nor the reducing end binding sites on the surface. This allows for the simplification:

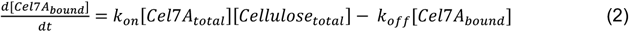

At steady-state, the time derivative goes to zero, hence:

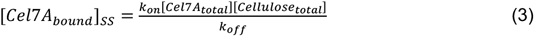

Using the initial condition of zero Cel7A bound at time zero, the solution to the differential equation is an exponential rise to the steady-state with rate constant *k*_*off*_(2):

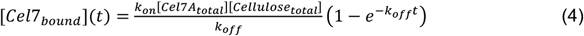

### Predicting rates of cellulose degradation in bulk assays

We can use the single-molecule results to predict expected results for cellobiose inhibition of Cel7A in bulk cellulose degradation assays. For approximating *k*_*cat*_, if we assume a 1 nm displacement per cellobiose released (3), then based on the velocity data in Fig. 3E, the *k*_*cat*_ can be modeled as:

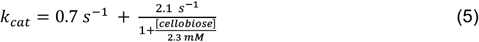

For approximating the *K*_*M*_, we can use the steady state Cel7A accumulation results from Fig. 2C, which gives a measure of the relative enzyme affinity for cellulose as a function of [cellobiose]. From Eq. 3,

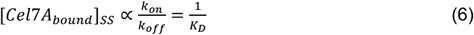

Hence, if we use the *K*_*D*_ as a proxy for the *K*_*M*_, then:

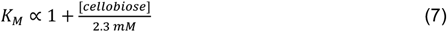

These results are plotted below.

**Figure SI 1:**
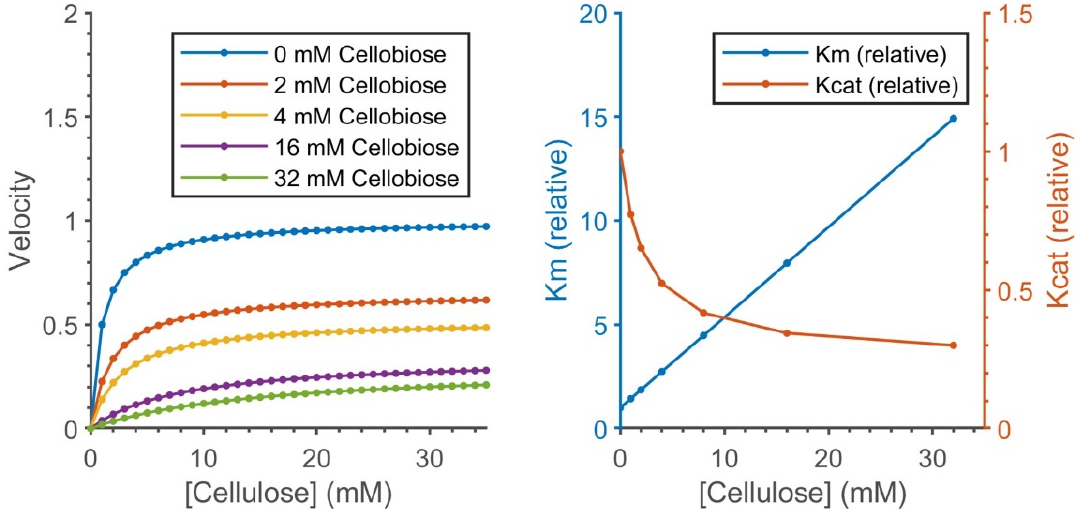
Cellulose degradation prediction in bulk assays under the presence of cellobiose. Maximum velocity (k_cat_) and K_M_ are both normalized to 1 under control (no cellobiose) conditions.

**Figure SI 2:**
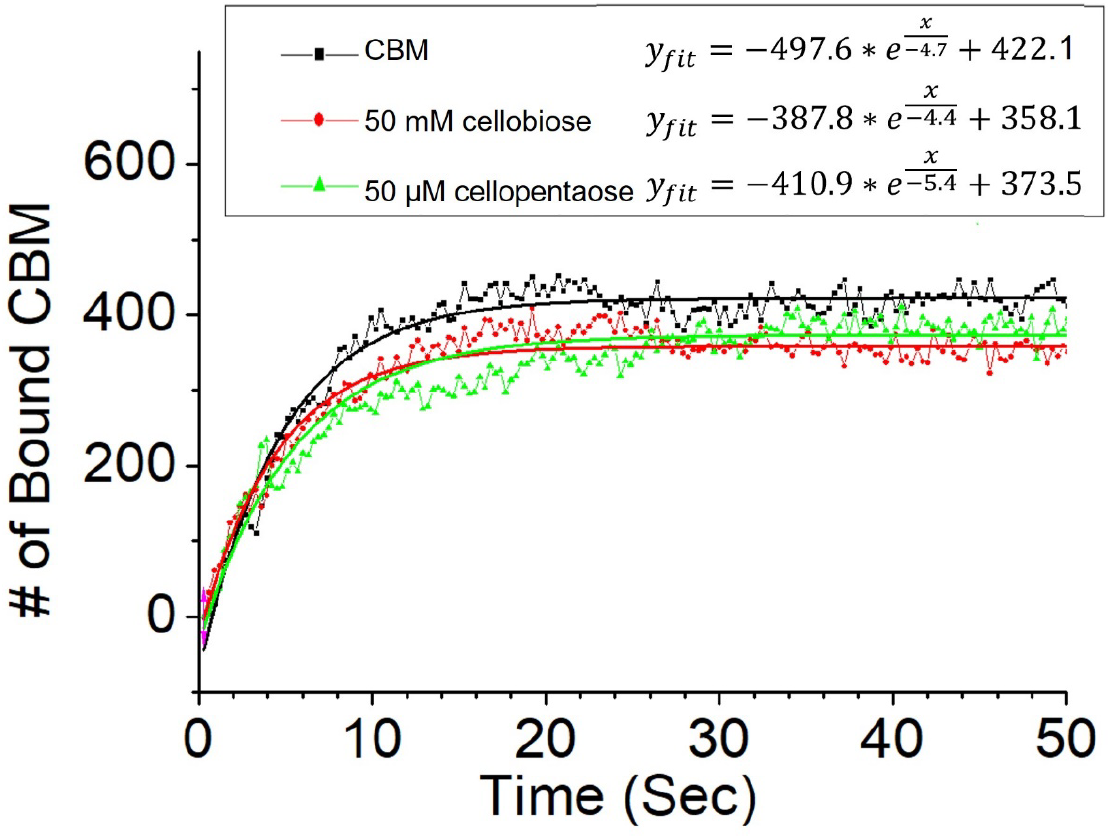
The binding of CBM3a to cellulose is unaffected by either cellobiose or cellopentaose.

